# Efficient Production of Pseudopregnant Female Mice Using Non-Estrous Individuals and the Whitten Effect

**DOI:** 10.1101/2025.02.06.636989

**Authors:** Yuji Noguchi, Eiji Watanabe

## Abstract

Pseudopregnant recipient females are essential for transplanting early mouse embryos during individualization. Traditionally, pseudopregnancy is induced through mating stimulation with vasectomized males, targeting females in proestrus or estrus. However, group housing of females in animal facilities often leads to fewer females entering estrus due to the Lee-Boot effect. This study focused on utilizing non-estrous females and the Whitten effect to produce pseudopregnant females. Females exhibiting characteristics of non-estrous states were visually selected and paired monogamously with vasectomized males for three consecutive days. This approach resulted in a high copulation success rate of 72.1% (CLEA Japan, Inc.) and 77.0% (Japan SLC, Inc.) on the expected third day. Subsequent embryo transfer using successfully copulated females resulted in offspring from 93.5% of ICR females from CLEA Japan, Inc. and 89.6% of ICR females from Japan SLC, Inc. Additionally, the suitability for embryo transfer could be determined by observing the condition of the ampulla of the oviduct. While this method requires a three-day preparation period compared to conventional approaches, it offers an effective alternative for creating pseudopregnant females for embryo transfer, potentially reducing animal use and associated costs.

## Introduction

When individualizing mouse embryos, recipient females with induced pseudopregnancy are required. Unlike many mammals, the corpus luteum of mice, which exhibits an incomplete estrous cycle, is usually in a dormant state and is activated by stimulation of the cervical canal. Such a state where the corpus luteum exhibits functionality is called pseudopregnancy [1, 2], which is generally induced by mating stimulation with a vasectomized male [3-8]. The conditions for planned mating of paired mice involve selecting females at the proestrus or estrus stage of their cycle [4, 6, 9]. The female’s estrous cycle, which repeats every 4-5 days, is divided into four stages: proestrus, estrus, metestrus, and diestrus [4, 6, 9-11]. Among these stages, only during estrus will the female accept the male and engage in mating [4, 6, 9]. Therefore, considering that the length of one estrous cycle in mice is typically 4-5 days, at least 4-5 times the number of females needed for embryo transfer will be required [5, 6]. On the evening before embryo transfer, through visual observation, females exhibiting characteristics such as “widely opened vaginal orifice with swollen, reddish-pink tissue [4, 5, 9, 11, 12] (Fig. 1A)” are identified. When these females are paired for infertile mating with vasectomized males, there is a high probability of copulation occurring that night [4, 9, 12]. The following day, successfully copulated females will have formed a whitish coagulated mass called a copulation plug in their external genitalia [1, 2, 6, 8, 9]. Females with observable copulation plugs are considered to have been induced into pseudopregnancy and are used for embryo transfer. The above-mentioned method has become widely established as a technique for generating pseudopregnant females today.

**Figure 1.**
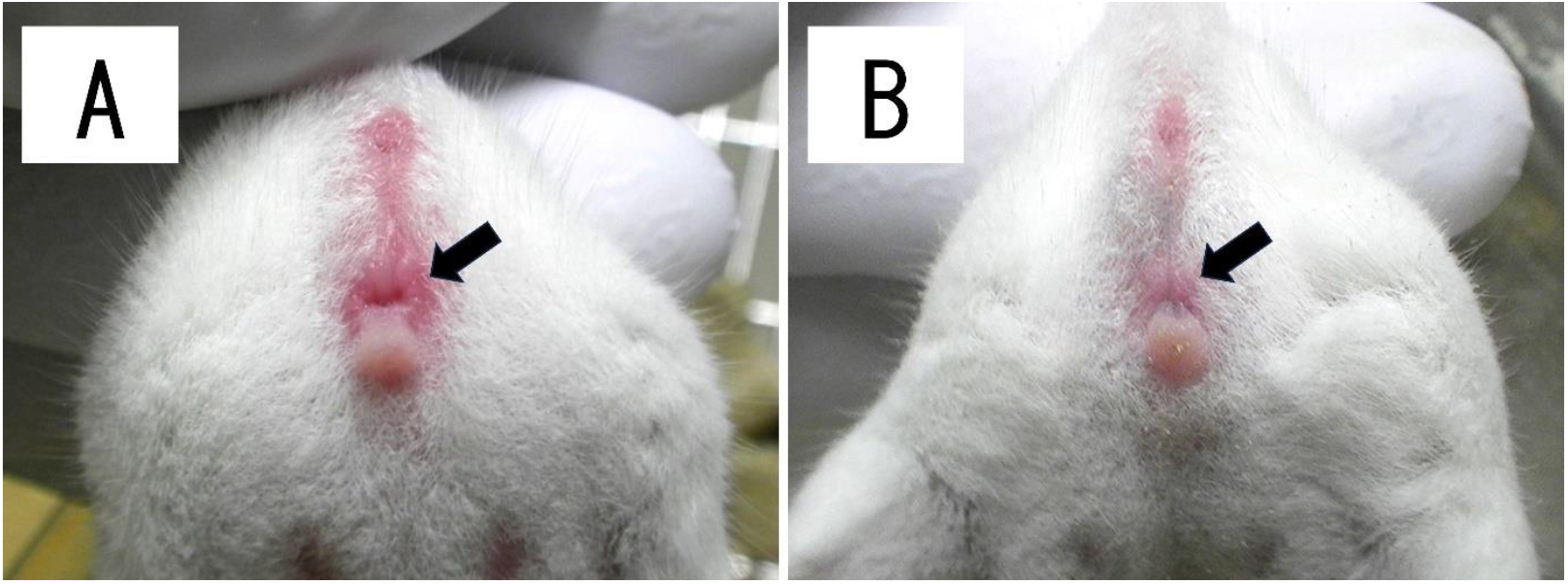
External genitalia of a female mouse (arrows). (A) External genitalia showing signs of estrus. When paired with a vasectomized male, there is a high probability of copulation occurring that night. These characteristics are key selection criteria for female mice in conventional pseudopregnancy induction protocols. (B) External genitalia in non-estrous state. These characteristics serve as selection criteria for female mice in mating protocols utilizing the Whitten effect. When paired with a vasectomized male, there is a high probability of copulation occurring on day 3.

However, mature females that typically come into heat every 4-5 days, when isolated from males and housed in group settings, may experience irregular estrous cycles or prolonged estrous cycles [13-16]. This phenomenon is known as the Lee-Boot effect. Among multiple volatile compounds excreted in urine via the adrenal glands [15, 17, 18], ‘2,5-dimethylpyrazine’ causes an effect of suppressing estrus [15, 18]. This female pheromone is contained in excreta and is not induced by visual or tactile stimuli [1, 19], but rather triggered by olfactory stimulation [11, 18] and mediated through the vomeronasal organ [20]. When females are housed in groups, exposure to ‘2,5-dimethylpyrazine’ excreted by other females leads to a loss of regularity in individual estrous cycles and a decrease in the frequency of estrus. Therefore, while there are strain differences, the estrus suppression effect becomes increasingly pronounced with higher population densities in group housing. When two females are housed together, the effect is minimal, but in small groups (4-8 mice), the diestrus phase is extended, resulting in a longer overall cycle [1, 18, 21, 22]. In larger groups (30 mice), estrus is completely suppressed [1, 18, 19]. In animal facilities that generate pseudopregnant females using conventional methods (selecting females during proestrus or estrus), these females are often maintained in group housing of approximately 10 individuals, primarily due to space constraints. As a result, various drawbacks arise due to the Lee-Boot effect. First, due to the prolongation of the estrous cycle, the frequency of estrus decreases, resulting in a reduction in the number of females in estrus [11, 22]. This makes it difficult to produce a consistent number of pseudopregnant females. Furthermore, the decreased frequency of estrus necessitates preparing more females, leading to a vicious cycle of increased housing costs, labor, and working hours. Additionally, maintaining a large female colony results in an increased proportion of mice that go unused and are ultimately discarded.

As a method to correct the Lee-Boot effect, the Whitten effect is well-known. The Whitten effect is a phenomenon where the presence of males improves the suppression of estrus in group-housed females [15, 23, 24]. When exposed to male urine, estrus becomes regular and synchronized, with most females progressing to copulation approximately 72 hours later [1, 11, 16, 19, 20, 22, 23]. This is because olfactory stimulation by various volatile components present in male urine induces estrus in females [15, 16, 22]. As this volatile component is an androgen-dependent pheromone in the air [11, 16, 25], it cannot induce the Whitten effect in castrated males [1, 16, 22, 26]. If the aforementioned Whitten effect could be effectively utilized, it might promote estrus that had been suppressed due to group housing, potentially allowing for more efficient production of pseudopregnant females. Therefore, in this study, we focus not on females during the conventional proestrus and estrus stages, but on females in a non-estrous state (late estrus or diestrus), and examined the production of pseudopregnant females through the Whitten effect. Although this method requires a three-day preparation period compared to conventional methods, it provides an effective alternative for creating pseudopregnant females for embryo transfer and has the potential to greatly reduce animal use and associated costs.

## Methods

### Animals and Rearing Environment

For the production of pseudopregnant females, ICR mice (ICR) were used, while C57BL/6J mice (B6) were used for collecting 2-cell stage embryos. ICR females were 10-19 weeks old, and B6 females were 4 weeks old. Males of both strains were 3-12 months old. These mice were obtained from CLEA Japan, Inc. (Tokyo, Japan) and Japan SLC, Inc. (Shizuoka, Japan). The animals were maintained under conditions of 22°C room temperature, 52% humidity, and a 12:12 hour light-dark cycle (lights on at 08:00), and provided with ad libitum access to standard rodent chow (CE-2; CLEA Japan, Inc., Tokyo, Japan) and water. Female ICR mice were housed in polycarbonate cages (CL-0105-2 Clean TPX, 263 × 422 × 150 mm; Japan CLEA Inc., Tokyo, Japan) in groups of 8-10 mice per cage after arrival from the breeder. After a 2-week acclimation period, they were used to produce pseudopregnant females starting at 10 weeks of age. The animal experiments described here were approved by the National Institutes of Natural Sciences Animal Experiment Committee (approval numbers: 21A005, 22A002, 23A028, 24A003) and conducted in accordance with the National Institutes of Natural Sciences Animal Experiment Regulations.

### Collection and Cryopreservation of Two-Cell Stage Embryos

In the evening (16:00-17:00), B6 females were intraperitoneally administered 0.1 ml of a superovulation-inducing agent [27] (CARD HyperOva®; Kyudo Co., Ltd., Saga, Japan). Forty-eight hours later, 5 IU of hCG (Gonatropin; Aska Pharmaceutical Co., Ltd., Tokyo, Japan) was administered intraperitoneally to induce superovulation.

In vitro fertilization (IVF) was performed 16 hours after the hCG injection. The IVF procedure was performed according to a previously described method [28] with slight modifications, using TYH medium (PHC Co., Tokyo, Japan). Before use, 0.2 ml of TYH medium was placed in a plastic dish (3.5 × 12 mm; Thermo Fisher Scientific K.K., Tokyo, Japan), covered with mineral oil, and equilibrated overnight in a 5% CO2, 37°C incubator to stabilize temperature and vapor phase. Oocytes were collected from the ampulla of the oviduct after superovulation. Sperm were obtained from the cauda epididymis of male B6 mice. After 90 minutes of pre-incubation of the sperm suspension in mineral oil-covered TYH medium, the sperm suspension was added to the 0.2 ml TYH medium containing the oocytes to achieve a final sperm concentration of 100-300 µl. After 3 hours of insemination, the oocytes were transferred to pre-equilibrated modified Whitten’s medium (mWM; PHC Co., Tokyo, Japan) [29] for further development. The following day, 2-cell stage embryos were used for embryo transfer or cryopreservation.

Cryopreservation of 2-cell stage embryos was performed using a simplified vitrification method [30]. SUMILON cryotubes (1.2 ml, MS-4501, Sumitomo Bakelite CO., Tokyo, Japan) were used. Thawing was carried out by adding 0.25 M sucrose (PHC Co., Ltd., Tokyo, Japan) warmed to 37°C, 30 seconds after transferring from liquid nitrogen to room temperature (23°C).

### Production of Pseudopregnant Females Using the Whitten Effect

During the morning hours (10:30-11:30), female ICR mice (10-19 weeks old, maximum 10 mice per cage) were selected for mating solely by visual observation of the external genitalia. Cytological examination of vaginal smears [4, 9, 11] was not performed because of the possibility that stimulation during cell collection from the vagina could affect the estrous cycle. The females were restrained by holding their tails and front legs against the cage lid. From the female group, mice showing characteristics of non-estrus (metestrus or diestrus) were selected, with the criteria of “absence of vulvar redness and swelling, and relatively small vaginal opening [4, 9, 11, 12] (Fig. 1B) “. These females were paired with vasectomized males for three consecutive days in a monogamous setting. On the morning of the final day (day 3), females with visible vaginal plugs were used as pseudopregnant females. The presence of vaginal plugs was checked every morning to confirm copulation success.

### Embryo Transfer

Morphologically normal 2-cell stage embryos, either fresh or frozen-thawed, were transferred to the oviducts of pseudopregnant females. For anesthesia, a mixture of medetomidine (0.75 mg/kg), midazolam (4 mg/kg), and butorphanol (5 mg/kg) was administered intraperitoneally (0.1 ml per 10 g body weight). Post-surgery, pseudopregnant females were administered a medetomidine antagonist (0.75 mg/kg) intraperitoneally to promote early recovery (0.1 ml per 10 g body weight). They were then kept warm on a 37°C heating plate and observed until they recovered from anesthesia and regained motor function. On day 20 or 21, when pseudopregnant females did not deliver naturally, viable offspring were recovered by cesarean section.

### Random Mating Control Group

Group-housed female ICR mice (approximately 20 weeks of age, maximum 10 mice/cage) were used. Since this was a test mating following vasectomy in males, females were randomly selected without any visual observation of external genitalia. They were continuously paired with vasectomized males from Monday to Friday. Mating was confirmed by checking for the presence of vaginal plugs each morning.

### Statistical Analysis

Statistical analysis was performed on data from the same period (December to May) to compare random mating and Whitten effect-based mating (CLEA Japan, Inc., Japan SLC, Inc.). The mating success rates on day 3 were analyzed using the chi-square test. A probability value of P<0.05 was considered statistically significant.

## Results

Three challenges are considered to be addressed when utilizing pseudopregnant females produced using the Whitten effect as recipient females in practical applications. These are: 1) whether they can be produced in a planned manner, 2) whether offspring can be obtained, and 3) whether the appropriate timing for embryo transfer can be determined. Each of these aspects was verified.

### Production of Pseudopregnant Females

While mating will eventually occur if males and females are housed together, unplanned production can interfere with experiments. Therefore, it is necessary to prepare pseudopregnant females to coincide with the embryo transfer date. In order to produce pseudopregnant females in a planned manner, a study was conducted to determine whether the estrus stage of the estrous cycle truly occurs on day 3 due to the Whitten effect. First, a total of 344 ICR female mice obtained from CLEA Japan, Inc. (Tokyo, Japan) were used in this study. As a result, 248 females (72.1%) showed vaginal plugs on day 3 (Table 1). Additionally, 9 females (2.6%) showed plugs on day 1, 41 (11.9%) on day 2, and 46 (13.4%) did not show any vaginal plugs (Table 1). In addition, 200 ICR female mice obtained from Japan SLC, Inc. (Shizuoka, Japan) were also used in this study. As a result, 154 females (77.0%) showed vaginal plugs on day 3 (Table 2). Additionally, 1 female (0.5%) showed plugs on day 1, 7 (3.5%) on day 2, and 38 (19.0%) did not show any vaginal plugs (Table 2). Vasectomized males, not being castrated, may have influenced the estrous cycle of females similarly to normal males [5]. Incidentally, in each instance, more than half of the paired mice successfully mated on the expected third day.

**Table 1.**
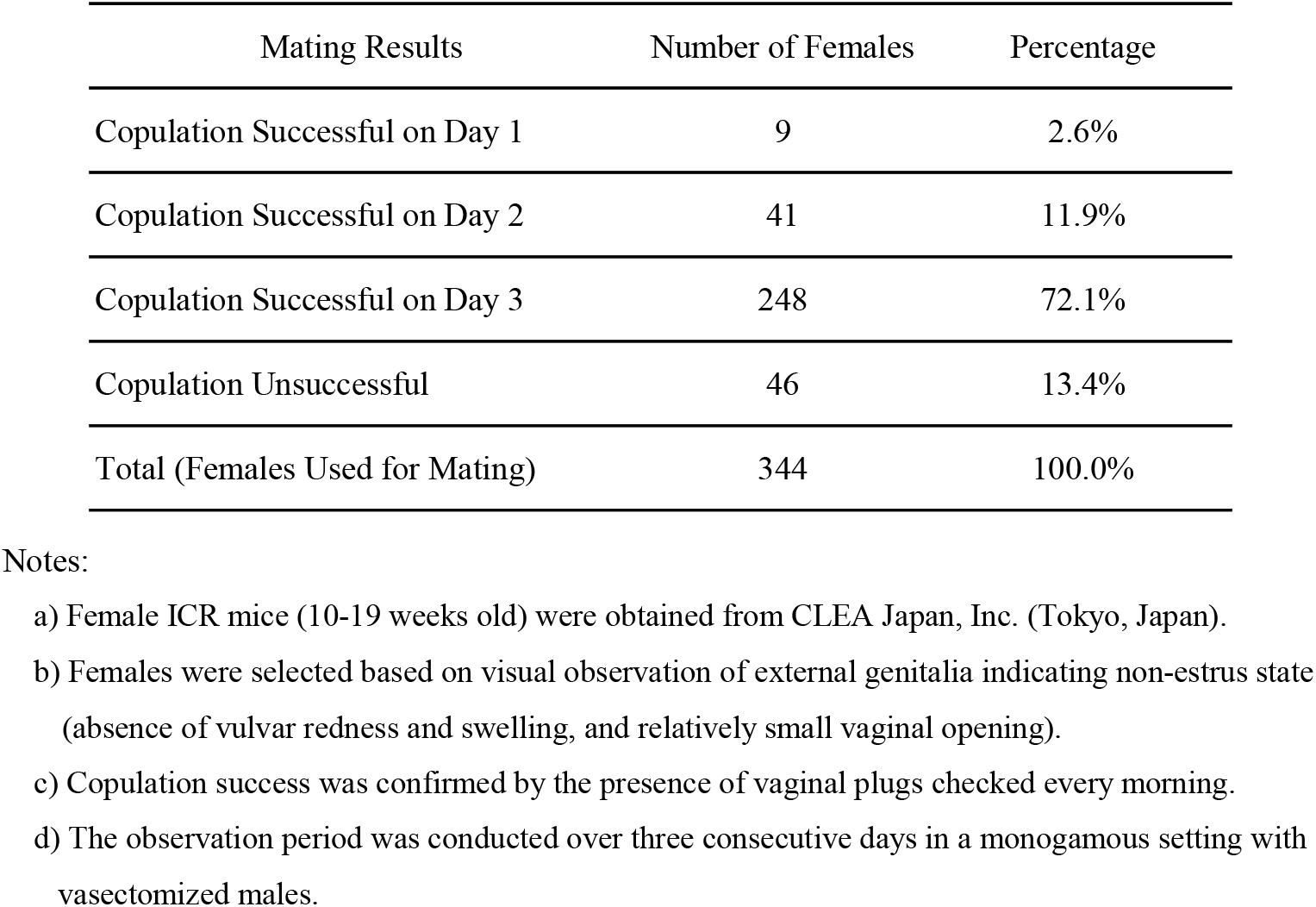
Results of Mating Using Non-Estrous Females (CLEA Japan, Inc.)

| Mating Results | Number of Females | Percentage |
| --- | --- | --- |
| Copulation Successful on Day 1 | 9 | 2.6% |
| Copulation Successful on Day 2 | 41 | 11.9% |
| Copulation Successful on Day 3 | 248 | 72.1% |
| Copulation Unsuccessful | 46 | 13.4% |
| Total (Females Used for Mating) | 344 | 100.0% |

Notes:
- a) Female ICR mice (10-19 weeks old) were obtained from CLEA Japan, Inc. (Tokyo, Japan). - b) Females were selected based on visual observation of external genitalia indicating non-estrus state (absence of vulvar redness and swelling, and relatively small vaginal opening). - c) Copulation success was confirmed by the presence of vaginal plugs checked every morning. - d) The observation period was conducted over three consecutive days in a monogamous setting with vasectomized males.

**Table 2.** Results of Mating Using Non-Estrous Females (Japan SLC, Inc.)

| Mating Results | Number of Females | Percentage |
| --- | --- | --- |
| Copulation Successful on Day 1 | 1 | 0.5% |
| Copulation Successful on Day 2 | 7 | 3.5% |
| Copulation Successful on Day 3 | 154 | 77.0% |
| Copulation Unsuccessful | 38 | 19.0% |
| Total (Females Used for Mating) | 200 | 100.0% |

Notes:
- a) Female ICR mice (10-19 weeks old) were obtained from Japan SLC, Inc. (Shizuoka, Japan). - b) Females were selected based on visual observation of external genitalia indicating non-estrus state (absence of vulvar redness and swelling, and relatively small vaginal opening). - c) Copulation success was confirmed by the presence of vaginal plugs checked every morning. - d) The observation period was conducted over three consecutive days in a monogamous setting with vasectomized males.

### Offspring Production

Of the 248 ICR female mice from CLEA Japan, Inc. that showed vaginal plugs on day 3, two-cell stage embryos (fresh and frozen-thawed) from B6 mice were transferred to the oviducts of 231 females (Table 3). As a result, 216 females (93.5%) produced offspring (including those obtained by cesarean section) (Table 3). Unfortunately, 15 females (6.5%) failed to establish pregnancy (Table 3). Similarly, of the 154 ICR female mice from Japan SLC, Inc. that showed vaginal plugs on day 3, two-cell stage embryos (fresh and frozen-thawed) from B6 mice were transferred to the oviducts of 115 females (Table 3). As a result, including those obtained by cesarean section, offspring were obtained from 103 (89.6%) of the 115 recipient females (Table 3). In contrast, pregnancy was not established in 12 (10.4%) recipient females (Table 3). In both groups of ICR female mice, embryo transfer was not performed when weak ampullary swelling was observed, due to the possibility that pseudopregnancy had not been induced. This occurred in 11 (4.4%) mice from CLEA Japan, Inc. and 9 (5.8%) mice from Japan SLC, Inc. (Table 3).

**Table 3.** Mouse Embryo Transfer Results Summary.

|  | Whitten Effect (CLEA Japan) |  | Whitten Effect (Japan SLC) |  |
| --- | --- | --- | --- | --- |
|  | Number of Females | Percentage | Number of Females | Percentage |
| **Total Embryo Transfers** | 231 | 93.1% | 115 | 74.7% |
| Successful Births (Including C-Sections) | 216 | 93.5% | 103 | 89.6% |
| Failed Pregnancies | 15 | 6.5% | 12 | 10.4% |
| **Not Transferred** | 17 | 6.9% | 39 | 25.3% |
| Canceled due to Insufficient Pseudopregnancy | 11 | 4.4% | 9 | 5.8% |
| Others | 6 | 2.4% | 30 | 19.5% |
| Total (Number of Mice with Vaginal Plugs) | 248 | 100.0% | 154 | 100.0% |

Notes:
- a) Embryos used: B6 strain mouse 2-cell stage embryos (fresh and frozen) - b) Transfer site: Oviduct - c) Pseudopregnancy assessment: Oviduct ampulla distension determines transfer eligibility - d) Embryo transfer performed on pseudopregnant females with clear ampullary swelling - e) Two different mouse suppliers: CLEA Japan, Inc. and Japan SLC, Inc.

### Timing determination for Embryo Transfer

When males and females cohabitate for three days, it becomes difficult to pinpoint exactly when mating occurred, making this determination crucial. This is because optimal timing for embryo transfer has been established in pseudopregnant recipients, including those prepared by the conventional method using proestrus females. For early-stage embryos, pseudopregnancy day 1 (0.5 dpc) is considered the appropriate timing. Indeed, embryo transfers performed on day 2 of pseudopregnancy rarely led to successful pregnancies. Fortunately, it was observed that the swelling of the ampulla of the oviduct changes daily, serving as a useful indicator. The ampulla of the oviduct exhibits marked distension on day 1 of pseudopregnancy (Fig. 2A) and undergoes significant shrinkage by day 2 (Fig. 2B).

**Figure 2.**
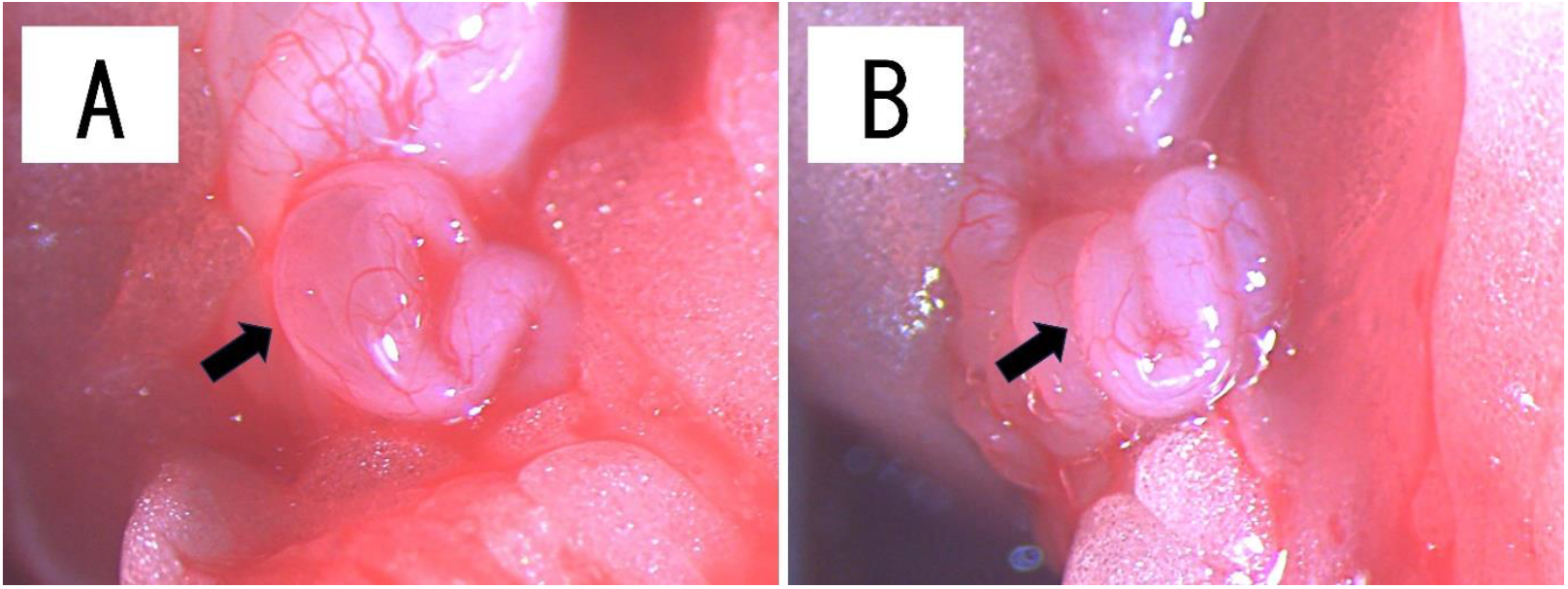
Ampulla of the oviduct in pseudopregnant female mice (arrows). (A) Day 1 of pseudopregnancy. The swollen ampulla indicates recent ovulation and estrus. Since pseudopregnancy is successfully induced by copulatory stimulation, this condition with the presence of a vaginal plug is considered optimal for embryo transfer. (B) Day 2 of pseudopregnancy. In contrast to day 1, no swelling of the ampulla is observed. Embryo transfer at this stage yielded poor results.

### Control Group with Random Mating

When group-housed females were randomly mated with vasectomized males, the highest copulation success rate was observed on day 3. However, not all matings were concentrated on day 3, with some occurring on other days as well (Table 4). Chi-square tests were performed to analyze the mating success rate on day 3, with successful copulation on day 3 defined as “mating success” and other outcomes as “mating failure.” The analysis revealed significant differences between random mating and both Whitten effect groups (Random vs CLEA Japan: χ^2^ = 4.32, df = 1, p < 0.05; Random vs Japan SLC: χ^2^ = 18.2, df = 1, p < 0.05). The copulation success rate on day 3 was significantly higher with Whitten effect-based mating compared to random mating, as females were selected through visual observation of external genitalia indicating non-estrus state.

**Table 4.** Comparison of Mating Success Between Random and Whitten Effect Mating (Dec.-May)

| Mating Results | A |  | B |  | C |  |
| --- | --- | --- | --- | --- | --- | --- |
|  | Random Mating |  | Whitten Effect (CLEA Japan) |  | Whitten Effect (Japan SLC) |  |
|  | Number of Females | Percentage | Number of Females | Percentage | Number of Females | Percentage |
| Copulation Successful on Day 1 | 17 | 16.8% | 5 | 3.5% | 1 | 1.1% |
| Copulation Successful on Day 2 | 9 | 8.9% | 19 | 13.5% | 4 | 4.4% |
| Copulation Successful on Day 3 | 54 | 53.5% | 94 | 66.7% | 75 | 82.4% |
| Copulation Successful on Day 4 | 7 | 6.9% | - | - | - | - |
| Copulation Unsuccessful | 14 | 13.9% | 23 | 16.3% | 11 | 12.1% |
| Total (Females Used for Mating | 101 | 100.0% | 141 | 100.0% | 91 | 100.0% |

Notes:
- a) Group A: Random mating (Monday-Friday); Groups B and C: Whitten effect-based mating (Tuesday-Friday) using females from CLEA Japan, Inc. and Japan SLC, Inc., respectively. - b) Values are expressed as number of females and percentages. - c) Hyphen (-) indicates no data collection on day 4 for Whitten effect groups.

## Discussion

In this study, the Whitten effect enabled high copulation success rates on day 3, consistent with expectations. The copulation success rates were 72.1% for ICR females from CLEA Japan, Inc. (Tokyo, Japan) and 77.0% for ICR females from Japan SLC, Inc. (Shizuoka, Japan). Subsequent embryo transfer using successfully copulated females resulted in offspring from 93.5% of ICR females from CLEA Japan, Inc. and 89.6% of ICR females from Japan SLC, Inc.

Moreover, we found that the feasibility of embryo transfer could be determined by observing the condition of the ampulla of the oviduct. Although this method produces females three days later than the conventional method using females in proestrus or estrus, these females can be effectively utilized as pseudopregnant recipients. By focusing on non-estrous individuals, the range of selectable females expanded, allowing us to obtain comparable results even with shorter working times and small-scale female stocks. Reducing the breeding scale may lead to a decrease in breeding costs, labor, and time. Furthermore, minimizing the number of females required to be maintained reduces unnecessary animal use, contributing to the 3R principles (Reduction, Refinement, Replacement) from an animal welfare perspective.

On the other hand, as evident from the results of the control group, not all pairings of group-housed females necessarily result in successful mating on the third day. The efficiency of producing pseudopregnant females may be somewhat compromised because non-estrous females are selected solely through visual observation, based on characteristics such as “a relatively small vaginal opening without redness or swelling of the external genitalia,” rather than performing cytological examinations of vaginal smears to determine each estrous cycle stage. In fact, it has been pointed out that even when selecting females in proestrus or estrus using conventional methods, the accuracy of observing external genitalia depends on the training and experience of the operator [4-6, 9]. However, the density of the female group and the duration of time spent in the group are positively correlated with the magnitude of the Lee-Boot effect [15, 31]. Prolonged group housing in the absence of males may enhance the Lee-Boot effect in females and potentially lead to more uniform responses to the Whitten effect [32]. Therefore, it is proposed that group-housing females for at least three weeks before male urine exposure [16, 20, 32] could stabilize the efficiency of producing pseudopregnant females by inducing estrus suppression in the majority of individuals.

Recently, the development of genome editing technology using the CRISPR/Cas9 system [33] has made the creation of genetically modified mice faster and easier. Against this background, it is suggested that the demand for pseudopregnant females has increased compared to before.

However, the production process has remained largely unchanged since the 1980s, and methods that are still criticized as inefficient continue to be employed [5, 6]]. To break through this situation, various methods for producing pseudopregnant females have been reported through the efforts of multiple researchers, and significant changes are emerging [4-8]. The Whitten effect has several advantages in producing pseudopregnant females: it is a simple and safe method requiring no mouse handling or injections, while being highly cost-effective [11, 32]. We hope this method will become one of the available options for producing pseudopregnant females, alongside various other newly developed methods.

## Acknowledgments

We would like to express our sincere gratitude to Dr. Tomoo Eto (Public Interest Incorporated Foundation Central Institute for Experimental Animals) for his valuable advice and insightful comments that greatly improved our manuscript. His expertise and guidance were instrumental in shaping this research. We also thank Mr. Daiji Fujimoto (formerly of Japan Animal Care Co., Ltd.) for his technical support.

